# Scale-dependent influences of distance and vegetation on the composition of aboveground and belowground tropical fungal communities

**DOI:** 10.1101/2020.06.01.127761

**Authors:** Andre Boraks, Gregory M. Plunkett, Thomas Doro, Frazer Alo, Chanel Sam, Marika Tuiwawa, Tamara Ticktin, Anthony S. Amend

## Abstract

Fungi provide essential ecosystem services and engage in a variety of symbiotic relationships with trees. In this study, we investigate the spatial relationship of trees and fungi at a community level. We characterized the spatial dynamics for above- and belowground fungi using a series of forest monitoring plots, at nested spatial scales, located in the tropical South Pacific. Fungal communities exhibited strong distance decay of similarity across our entire sampling range (3–110,000 m), and also at small spatial scales (< 50 m). Unexpectedly, this pattern was inverted at an intermediate scale (3.7–26 km). At large scales (80–110 km), belowground and aboveground fungal communities responded inversely to increasing geographic distance. *Aboveground fungal community* turnover (beta diversity) was best explained, at all scales, by geographic distance. In contrast, *belowground fungal community* turnover was best explained by geographic distance at small scales, and tree community composition at large scales. We demonstrate scale-dependent spatial dynamics of fungal communities, synchronous spatial dynamics for trees and fungi, and the varying influence of habitat versus geographic distance in structuring Soil, *Selaginella sp.,* and Understory fungal communities.

## Introduction

Fungi and plants have coexisted for millions of years, often forming important symbiotic relationships. Fungi structure forest communities [1, 2] both through antagonism (by causing negative density-dependent growth and mortality [3]) and through mutualisms (as with mycorrhizae [4]). Similarly, plants impact fungal community composition through environmental modification or specificity [5]. Whilst co-occurrence patterns of bipartite relationships between host plants and fungi are important for understanding population-level dynamics, a more complete picture of spatial dynamics in complex ecosystems might be drawn at the community level. Estimates of fungal diversity range from 1.5 to 3.8 million species [6]. Plant species richness is predicted to be around 450,000 species, two thirds of which are found in the tropics [7]. A framework for examining interactions between co-occurring communities is to compare their compositional turnover along gradients.

The distance decay of similarity (DDS) is a recurrent phenomenon describing how the relationship between two entities changes over geographic space, a pattern consistent with Tobler’s first law of geography [14]. Tobler’s law intuitively states that nearby things have a tendency to be more similar than distant things. Within community ecology, DDS describes a pattern of community-membership turnover (beta-diversity) with increasing geographic distance and is used to uncover patterns of species distribution and aggregation [15]. The distance decay relationship is a powerful tool in spatial ecology because the slope of this relationship reflects a combination of environmental and biotic variables [16]. Ecologists have used the DDS to infer the relative importance of such divers topics as dispersal limitation in trees [17] and in the niche partitioning of diatoms [13]. For communities of bacteria, in general, there is recurrent evidence of a distance-decay relationship predicting community-composition divergence positively correlated with geographic distance [18–20]. Distance decay of similarity is not a constant in community ecology [21, 22] which raises questions about the circumstances that lead exceptions of distance-decay relationships [23].

Distance-decay relationships are both scale and system dependent, so recognizing scale-dependency is an important step to revealing insights into processes driving patterns of biodiversity. Scale-dependent patterns were documented in microbes, indicating that the relative importance of mechanisms generating spatial structure, such as dispersal limitation and environmental filtering, vary by geographic scale [24]. Aboveground, the fungal phyllosphere community may [25, 26] or may not [27] exhibit scale-dependent DDS relationships, even showing variable results within the same study [28]. Belowground, soil fungi exhibit distance decay relationships that vary by soil horizon [29]. Spatial structure among ectomycorrhizal communities is related to host density at a local scale, but climate seems to be more important at a global scale [30]. The relative importance of mechanisms generating spatial structure, such as dispersal limitation and environmental filtering, are scale and habitat dependent. Most studies of spatial processes address an individual habitat type or scale, making synthesis of results across studies difficult due to the high variability in sampling methodology [31]. There is a clear need for the consideration of multiple habitats and spatial scales, simultaneously, to allow for an ecosystem-wide perspective within complex forest systems.

Characterization of the spatial dynamics for both fungi and plants from the same environment provides us with a critical window into the ecology of complex forest systems [8]. Plant communities have been studied for centuries under the lens of spatial ecology [9]. By comparison, descriptions of the distribution of fungal communities remain relatively less common. Hawksworth & Lücking [6] estimated less than 10 percent of all fungal species have been formally described, leaving our understanding of fundamental ecological tenants for fungi behind those of animals and plants. Recent studies have highlighted how little we know of the natural distribution patterns of fungi [10, 11], and ongoing research has yet to establish the extent to which fungi and plants exhibit similar biogeographic patterns [12, 13]. We now have the capacity to simultaneously investigate biogeographic patterns of fungi and plants by combining high-throughput sequencing technology and long-term vegetation monitoring transects.

In this study, we describe the spatial distribution patterns of fungal communities from multiple habitat types (Soil, Understory, and *Selaginella)* and multiple spatial scales. We then integrated plant community data, collected in parallel with the fungal data, to link distribution patterns of trees and fungi. The aim of this study was to assess whether fungal community beta-diversity, derived from three different habitats, vary over three geographical scales [ranging from local (3.33–37.23 m), to within islands (3.7–26 km), to between islands (80–110 km)], and to what extent fungal spatial dynamics are synchronous with tree communities. We characterized the tree and fungal communities from tropical forests found on the nearby islands of Aneityum and Tanna, in the South Pacific archipelago of Vanuatu, to address the following questions: (i) do fungal communities show distance-decay patterns at multiple geographic scales? If so, (ii) does the strength of decay vary with the geographic scale of investigation? And, (iii) to what extent is fungal community beta-diversity attributable to patterns in plant-community diversity and distribution? We predicted that the rate of community turnover would increase with geographic scale, and that these scale-dependent relationships would vary dependent on whether the fungal community was sampled from either Soil, Understory, or *Selaginella.* Further, we expected a colinear relationship between plant and fungal beta-diversity, such that each would display similar patterns of correlated scale-dependent community turnover. Evidence of scale-dependent patterns among plants and fungal communities provide us with new insights into the factors governing biodiversity in tropical forests.

## Materials and Methods

### Study site and sampling

The Republic of Vanuatu is an archipelago of more than 80 islands located in the Southwestern Pacific. Sampling for this project occurred in the southernmost province of Tafea, on the islands of Tanna and Aneityum (Figure 1). Recent efforts to document the flora and vegetation of Tafea have let to a network of sampling transects that were used in this study. Sampling efforts on Tanna and Aneityum primarily occurred in habitats typified as low to mid-elevation rain forest [33]. Fieldwork occurred over the span of two trips during August 2017 (Aneityum) and December 2017 (Tanna). The two islands are separated by 86 kilometers of open ocean.

**Fig. 1.**
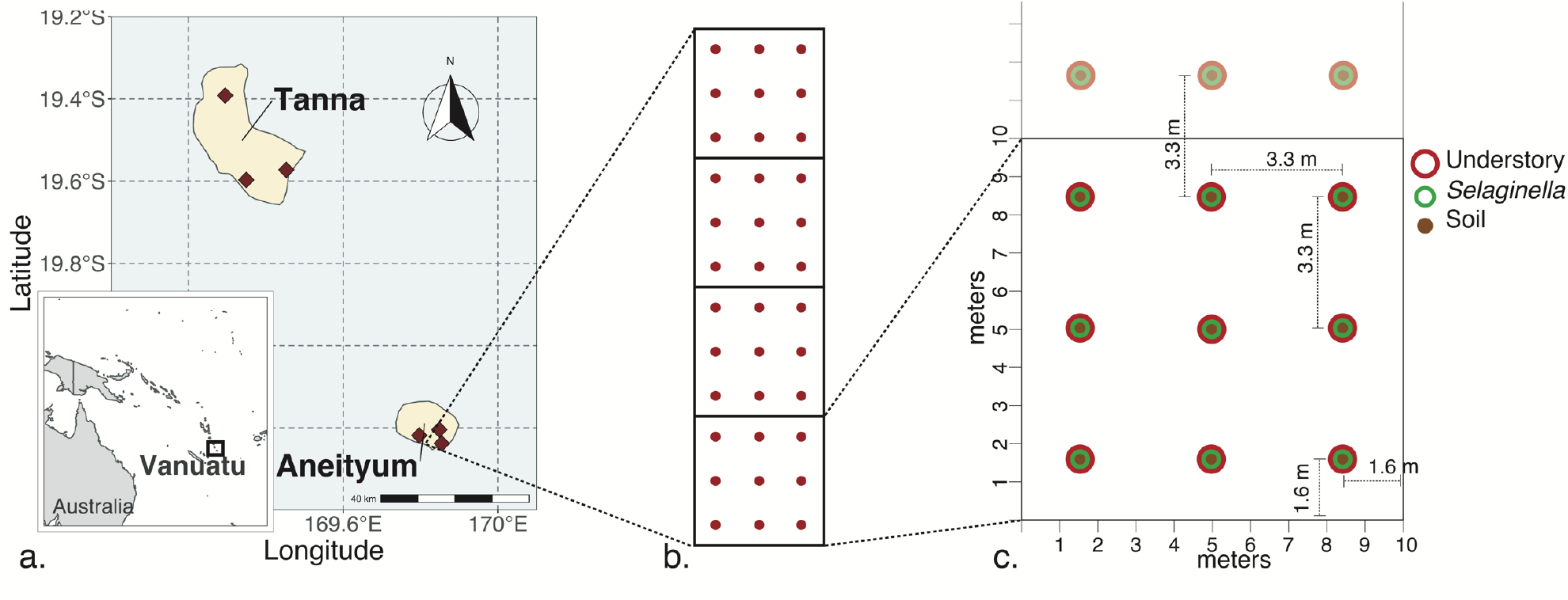
Location of the 6 transects sampled in this study. Three transects were each sampled on the islands of Tanna and Aneityum, Vanuatu (a). Each transect contained 36 sampling sites, split equidistant among 4 contiguous plots (b). At each sampling site one fungal community was harvested from each of three habitats; Soil, Selaginella, and Understory (c, Online Resource 1).

Three long-term vegetation monitoring transects were sampled on each island. The average distances between transects within islands is 5.45 Km (Aneityum) and 21.94 Km (Tanna) (Figure 1a). Transects dimensions were 10 m by 40 m. The vegetation in each transect was catalogued and all trees greater than 5 cm (diameter at breast height) were mapped and identified. Although plant species differed among plots, *Selaginella* dominated the understory throughout, which is why we targeted this species for sampling.

To characterize the fungal communities occurring within the transects, 36 sampling sites were established in a grid within each transect (Figure 1b). At each sample site, three separate samples were taken: Soil, *Selaginella* sp. (Lycopodiaceae)), and Understory (swabbed angiosperm leaf surface) (Figure 1c). The details of sampling methodology for each sample source is outlined in Online Resource 1. *Selaginella* voucher specimens have been accessioned in the Joseph Rock Herbarium at the University of Hawai’i. In total, 6 transects, each containing 36 sampling sites, at which there were 3 sampling events (Soil, *Selaginella,* and Understory), resulted in 648 fungal community samples. Samples were transported to the laboratory at the University of Hawai’i at Mānoa. Desiccated *Selaginella* samples were pulverized in a biosafety cabinet using sterile mortars and pestles and liquid nitrogen. Both *Selaginella* powder and CTAB swabs were stored at −20 °C until DNA extraction.

### Fungal DNA extraction, PCR amplification, and sequencing

DNA extraction was performed using the Qiagen DNeasy PowerSoil DNA isolation Kit (Qiagen, Cat No./ID: 12888). All extraction steps were performed in a biosafety cabinet to minimize environmental contamination. Negative control DNA extractions were run using sterile swabs and CTAB solution that traveled to the field but did not come in contact with leaf material.

Amplicon libraries were prepared in a single PCR reaction using Illumina-barcoded fungal-specific primers. Primers ITS1F and ITS2 were used to target the hypervariable nuclear ribosomal ITS1 region which is flanked by the 18S and 5.8S nrDNA regions. The ITS1 region was amplified using 8-base-pair indexed primers (0.2 μM), gDNA (~ 5 ng) and Phusion Master Mix (New England Biolabs, Massachusetts), using thermocycler parameters recommended by <u>Phusion</u>. Fungal libraries were purified and normalized using Just-a-plate Purification and Normalization Kit (Charm Biotech, San Diego, California) and quantified on a Qubit fluorometer (Invitrogen, Carlsbad, California). Samples were pooled and submitted for sequencing at the Genomics Core Facility at the University of California–Riverside, Institute for Integrative Genome Biology. The Core Facility conducted quality control using a Bioanalyzer and qPCR to optimize cluster density. Sequencing occurred on the Illumina MiSeq platform (NextSeq500 Sequencer) using V3 chemistry (Illumina Inc., San Diego, CA), allowing for 300 bp paired-end reads.

ITS1 sequences were extracted from the flanking ribosomal subunit genes using *ITSxpress* (Rivers et al., 2018), then filtered by quality scores using the *FASTX-Toolkit* (Hannon, 2010). Reverse reads were discarded due to low quality reads. Chimeras were detected and removed using *vsearch* (Rognes et al., 2016). Sequence were clustered at 97% identity and fungal taxonomy was assigned using the Python package *constax* (Gdanetz et al., 2017) and the unite database, via consensus of three separate taxonomy classification algorithms. Putative contaminants were identified based on their prevalence in extraction and PCR negative controls and removed using the R package *decontam* (Davis et al., 2018). OTUs that could not be assigned to a fungal phylum were discarded. Differences in sampling depth between samples were normalized using a variance stabilizing transformation (VST) [35] in *DESeq2* [36] within R [37]. All downstream analyses were performed in R and used the VST transformed dataset except in cases when measurements of alpha diversity were employed, in which case the OTU table was rarefied to a minimum sequence depth of 3,000 sequence reads. OTU read abundance data, taxonomic assignments, sample metadata, and ancillary collection data were compiled with the R package *phyloseq* (McMurdie & Holmes, 2013).

### Data analysis

Spatial autocorrelation for tree and fungal communities were assessed using Mantel tests. Bray-Curtis dissimilarity matrices were calculated for Hellinger-transformed fungal communities using *vegan*::vegdist. Geographic distance was calculated using *fossil*::earth.dist. The decay slope at each spatial scale was calculated using a linear regression and significance was tested by permutation. Distance decay rates were calculated for the entire study and at three subset spatial scales: within transects (3.33–37.23 m), between transects within islands (3.7–26 km), and between islands (80–110 km). The minimum grain size for fungal and tree communities differed in that fungal communities can be represented at the grain size of the individual sample site (i.e., one of the 36 per transect), but tree community grain size could not be reduced beyond that of the transect level. For this reason, we were able to calculate distance decay within transects for the fungal dataset, but not the tree dataset. To test whether fungal distance decay relationships were related to tree community distance decay relationships, we performed partial Mantel tests on Bray-Curtis tree community dissimilarity and Bray-Curtis fungal community dissimilarity while accounting for any collinearity associated with geographic distance.

Variation in fungal communities as a function of geographic distance and tree community was tested using Generalized Dissimilarity Modeling (GDM; R package *gdm*) [39]. GDM provides a non-linear perspective on the relative importance of geographic distance and tree community in structuring fungal communities at various scales. GDM coefficients (the maximum height of its spline [39, 41]) can be interpreted as the amount of variation explained by a predictor variable when all other model variables are held constant. Variation in the GDM spline slope is also informative of how the model varies over a range. Significance of each GDM variable was tested by permutation. We tested how dissimilarity of sample OTU composition between transects varied with differences in tree community and geographic distance (fungal community ~ tree community + geographic distance).

## Results

### Fungal Species Identification through Sequencing

Despite efforts to maintain a balanced sampling design, not all sampling sites contained *Selaginella* growing at the pre-determined location and not all soil or phylloplane (understory epiphyte) samples resulted in sequence data. The final number of fungal metagenome samples totaled 580 (Soil = 216, Understory = 213 and *Selaginella* = 151). Within these samples we identified a total of 27,581 OTUs (clustered at 97 % similarity) of which 10 OTUs were identified as contaminants and were removed from the dataset. We then culled OTUs that could not be identified as Fungal at the phylum level reducing our dataset to 18,147 were identified as Fungal at the Phylum taxonomic rank and were retained for downstream analysis. Each sample type varied significantly in the number of OTUs (Tukey’s test;*p* < 0.01). Understory had the largest average number of OTUs 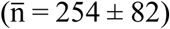, followed by *Selaginella* 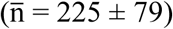, and Soil 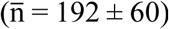. Alpha diversity of fungi (measured as total number of observed OTUs) also varied by plot, transect, and island (Online Resource 2).

### Spatial dynamics of fungal and tree communities

Mantel tests showed significant distance-decay patterns. In general, as geographic distance increased, fungal communities became increasingly dissimilar for all sample types (Figure 2). This general trend did not hold true when sub-setting among various spatial scales. At the smallest geographic scale, within transects(< 40 m), all fungal communities follow a distance decay pattern similar to the general trend seen across the extent of the study. At an intermediate scale, between transects (3.7–26 km), fungal communities invert the general trend of distance decay, such that fungal-community composition increases in similarity with growing spatial distance. At the largest scale, between islands (80–110 km), fungal communities vary in distance decay response. *Selaginella* fungal communities decreased in similarity with distance, whereas Soil fungal communities increased in similarity with distance. Linear regressions fit to pairwise sample distances indicated significant scale-dependent patterns present in all fungal communities at each scale, except for understory fungi between islands (80–110 km; Table 1). Tree communities (Mantel *r* = 0.357,*p* < 0.001) exhibited a distance decay slope across the extent of the study (0–110 km) and was similar to what was observed among the fungal communities (Figure 2; Table 1).

**Fig. 2.**
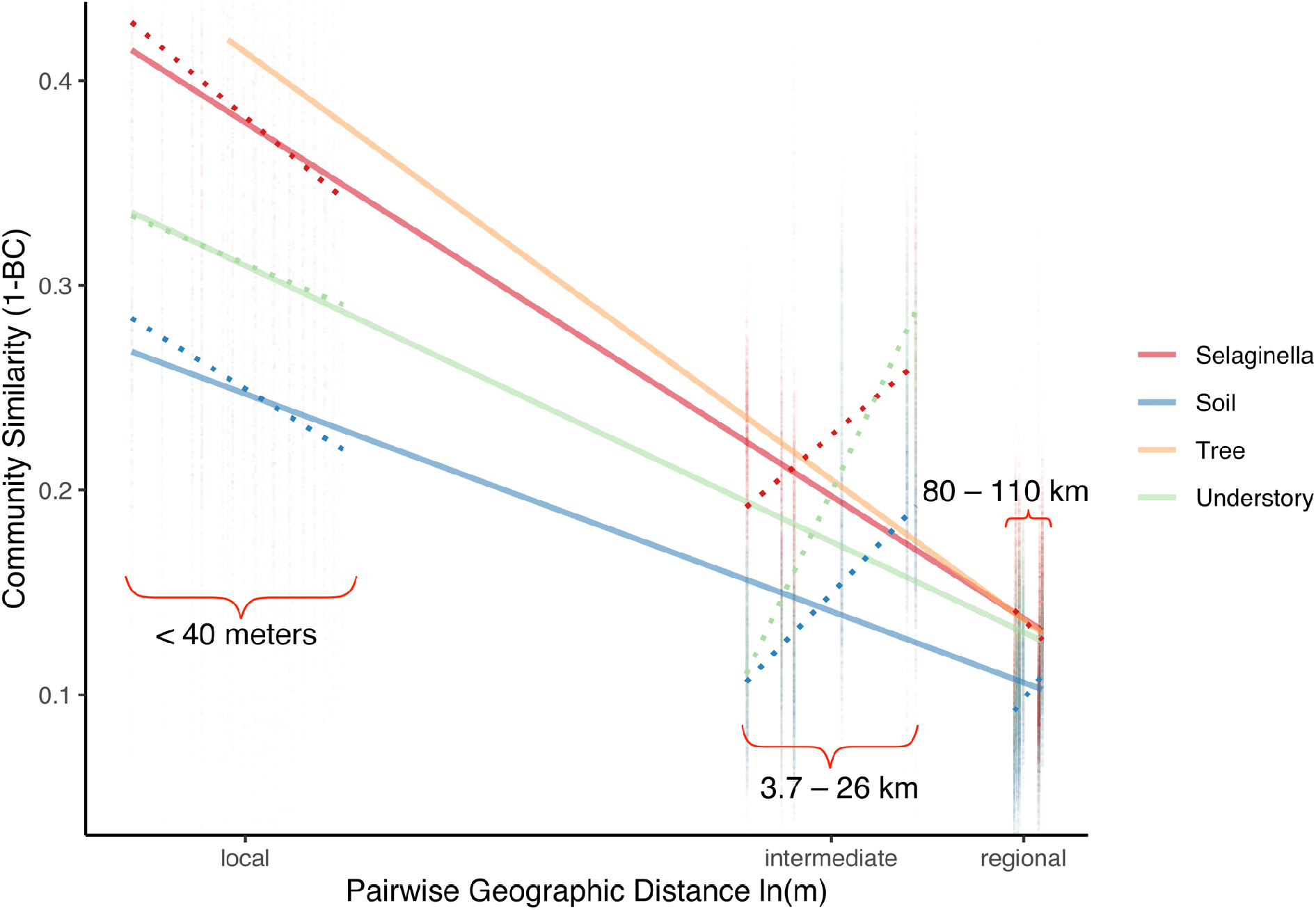
Distance decay of similarity (DDS) relationships for fungal (Soil, Selaginella, Understory) and tree communities across various scales. In general, fungal communities become less similar with growing isolation. For fungal communities, there is an inversion of this trend occurring at intermediate scales. Regression lines denote the leastsquares linear regression across the extent of the study. The three scales tested in this study are indicated with dotted lines and curly brackets: local (3.33–37.23 m), across islands (3.7–26 km), between islands (80–110 km). Geographic distances are the natural log of meters between sample sites. Dissimilarity between communities is calculated as the natural log of Bray-Curtis dissimilarity. Only the regression lines for significant Mantel tests (p < 0.05) are shown in this figure. Non-significant Mantel test results can be found in Table 1. Similarity measured as 1-Bray-Curtis dissimilarity.

**Table 1.**
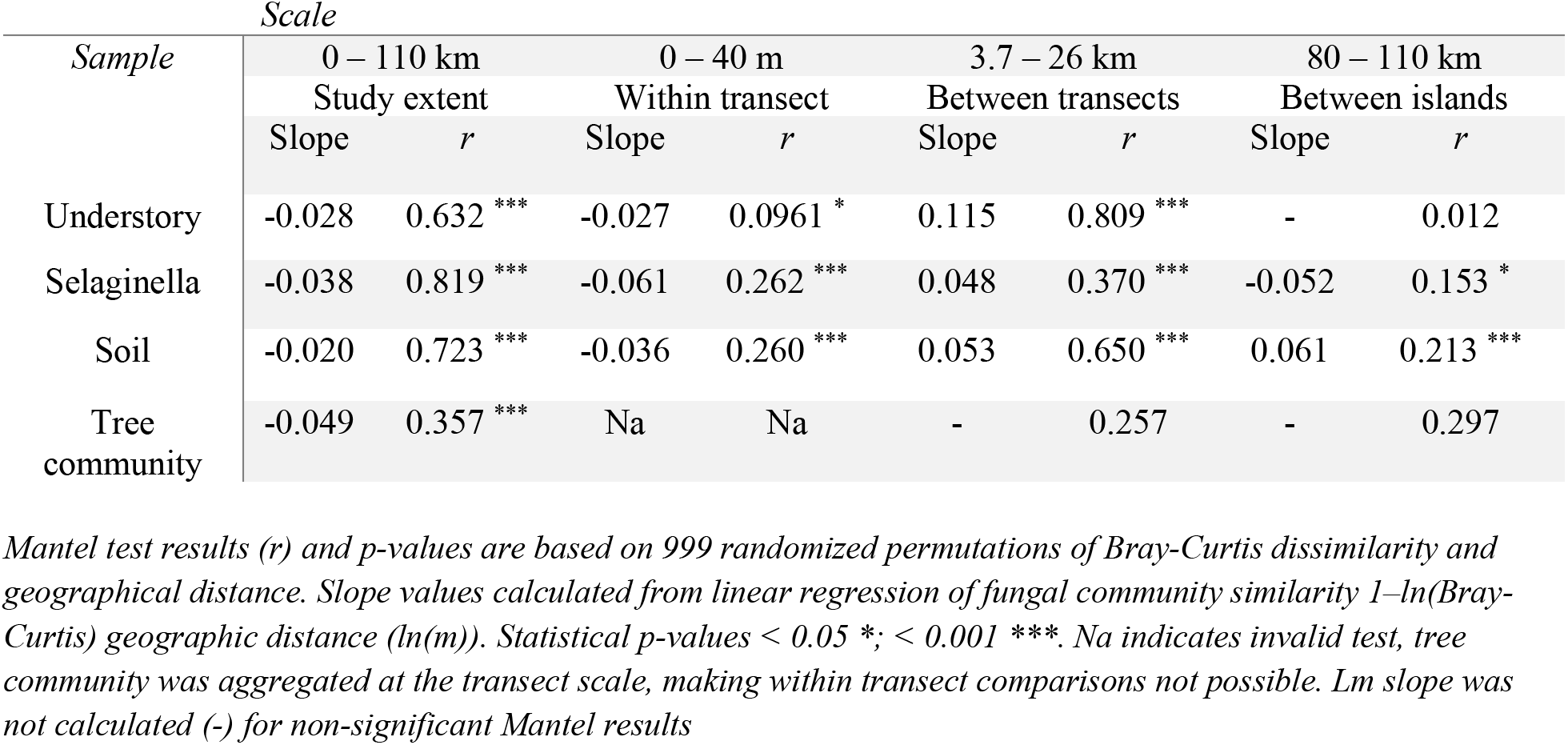
Mantel test summary statistics for decay of fungal and tree community similarity with geographic distance. Fungal communities were derived Soil, Selaginella, and Understory foliar epiphytes) spanning various geographic scales.

| Sample | Scale |  |  |  |  |  |  |  |
| --- | --- | --- | --- | --- | --- | --- | --- | --- |
|  | 0 – 110 km |  | 0 – 40 m |  | 3.7 – 26 km |  | 80 – 110 km |  |
|  | Study extent |  | Within transect |  | Between transects |  | Between islands |  |
|  | Slope | *r* | Slope | *r* | Slope | *r* | Slope | *r* |
| Understory | -0.028 | 0.632 *** | -0.027 | 0.0961 * | 0.115 | 0.809 *** | - | 0.012 |
| Selaginella | -0.038 | 0.819 *** | -0.061 | 0.262 *** | 0.048 | 0.370 *** | -0.052 | 0.153 * |
| Soil | -0.020 | 0.723 *** | -0.036 | 0.260 *** | 0.053 | 0.650 *** | 0.061 | 0.213 *** |
| Tree community | -0.049 | 0.357 *** | Na | Na | - | 0.257 | - | 0.297 |

Mantel test results (*r*) and *p*-values are based on 999 randomized permutations of Bray-Curtis dissimilarity and geographical distance. Slope values calculated from linear regression of fungal community similarity $1 - \ln(\text{Bray-Curtis})$ geographic distance ( $\ln(m)$ ). Statistical *p*-values < 0.05 \*, < 0.001 \*\*\*. Na indicates invalid test, tree community was aggregated at the transect scale, making within transect comparisons not possible. Lm slope was not calculated (-) for non-significant Mantel results

### The relationship between tree and fungi community composition

The relationship between tree community and fungal community composition was examined using Mantel tests, partial Mantel tests (Table 2), and GDM. Mantel tests showed a positive correlation between plant and fungal community Bray-Curtis dissimilarity, indicating synchronous beta-diversity turnover. As plant community became dissimilar, so too did the fungal community for all three sample types (Soil r = 0.612, Selaginella r = 0.632, Understory r = 0.556; *p* = 0.001). This relationship held true when accounting for the collinearity of geographic distance using partial Mantel tests (fungal community x plant community x geographic distance) (Soil r = 0.535, Selaginella r = 0.548, Understory r = 0.452; *p* = 0.001) (Online Resource 3).

**Table 2.** Mantel correlation statistics (r) and P-values between tree community composition and fungal community composition dissimilarities (Bray–Curtis) for three fungal sample sources: Soil, Selaginella, and Understory. Partial Mantel correlations, comparing tree and fungal community composition, while accounting for pairwise geographic distance collinearity. This table indicates that tree community composition is associated with a particular fungal community composition. Tree community compositional turnover is correlated with fungal community turnover.. The partial Mantel test demonstrates how this community-community decay pattern is present irrespective of geographic distance.

| Fungal community | Tree community |  |
| --- | --- | --- |
|  | Mantel's *r* | *p* |
| Understory | 0.556 | 0.001 |
| Selaginella | 0.632 | 0.001 |
| Soil | 0.612 | 0.001 |
| Partial Mantel: controlling for geographic distance |  |  |
| Understory | 0.452 | 0.001 |
| Selaginella | 0.549 | 0.001 |
| Soil | 0.535 | 0.001 |

GDM models testing the relative importance of both geographic space and tree community composition on structuring aboveground or belowground fungal community composition (fungal community ~ tree community + geographic distance) indicated that geographic distance, rather than tree community, better explained compositional dissimilarity for fungal communities aboveground; *Selaginella* (GDM coefficient; geography = 1.023, tree community = 0.301) and Understory (GDM coefficient; geography = 0.682, tree community = 0.634) (Figure 3; Understory, *Selaginella).* By contrast, tree community, rather than geographic distance, explained more of the compositional dissimilarity for fungal communities belowground; Soil (GDM coefficient; geography = 0.550, tree community = 0.721) (Figure 3; Soil).

**Fig. 3.**
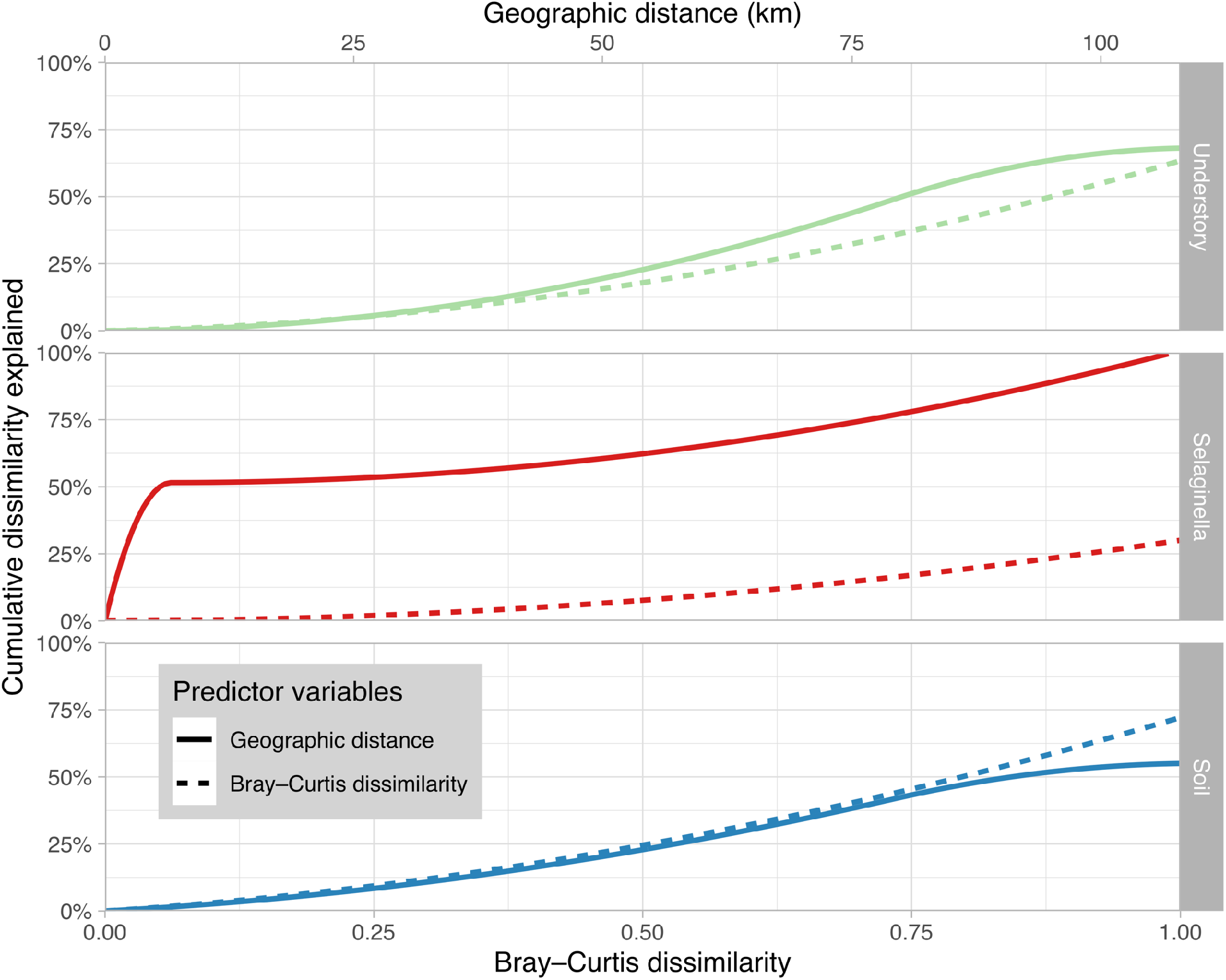
Splines for generalized dissimilarity models (fungal community ~ tree community + geographic distance). Models quantify the relationship between tree community (dotted line) and geographic isolation (solid line) and their effect on structuring fungal communities of Selaginella, Soil, and Understory. The maximum height of a spline can be interpreted as the total contribution that a factor explains of the observed differences in fungal communities when all other variables in the model are held constant. The slopes of the GDM spines indicate the relative importance for a predictive variable across its range. Geographic distance best predicts fungal communities of Selaginella and Understory across the entity study, whereas, tree community is a better predictor of Soil fungal communities at large distances (>75 km).

The slopes of the GDM splines also indicate non-linear relative importance of a predictive variable across its range. For example, the slope of geographic distance spline explaining soil fungi is steepest at intermediate ranges (Figure 3; Soil), indicating that geographic distance is most explanatory of soil fungal communities at an intermediate range (~ 30–80 km). By comparison, geographic distances around 100 km explain little of the variation observed in soil fungal community composition (Figure 3; Soil).

## Discussion

For biological communities, the slope of a distance-decay relationship is a function of both the environmental factors that act upon the community (exogenous factors) and organismal characteristics (endogenous factors) [16]. Our results show significant variability in the distance decay relationship of tropical fungal communities and indicate that this variability is dependent on both scale and habitat. At the local scale (< 45 m), above- and belowground fungal communities show significant distance decay. At intermediate distances (3.7–26 km), fungal communities invert distance decay relationships so that, as geographic distance increases fungal communities become more similar. Distance decay patterns at the regional scale (~ 80–100 km) are mixed with negative spatial autocorrelation belowground and positive spatial autocorrelation aboveground (Figure 2). The observed variation in distance-decay relationships indicates that the relative influence of environmental and organismal characteristics on community dynamics is system and scale dependent. These results are corroborated with our GDM model. Fungal communities are strongly influenced by geographic distance at local and intermediate scales. At large scales (~100 km), vegetation composition is more important in structuring belowground, but not aboveground fungal communities.

Identifying how dispersal and environment independently structure community composition is difficult in microbial systems. In this study we observed positive spatial autocorrelation at short distances of less than 50 meters, indicating geographic distance between samples was a strong predictor of community similarity. Positive spatial autocorrelation is representative of an aggregating spatial dynamic (Figure 4a), whereby fungal communities are clustered at a local scale. Similar results have been observed in other studies [30] looking at spatial autocorrelation of soil fungal communities over short distances of < 10 m [42], < 50 m [29], and < 1 km [25]. Dispersal limitation is an important factor at small scales [43], although we are unable to determine its specific contributions to our results. Accurately measuring the dispersal distance of fungal propagules remains a complicated task. Fungi likely have a longtailed dispersal distribution, whereby the vast majority of reproductive propagules travel over a distance of centimeters and meters, rather than kilometers, a process that would generate the fungal community aggregation patterns observed at small scales in Vanuatu (Figure 2).

**Fig. 4.**
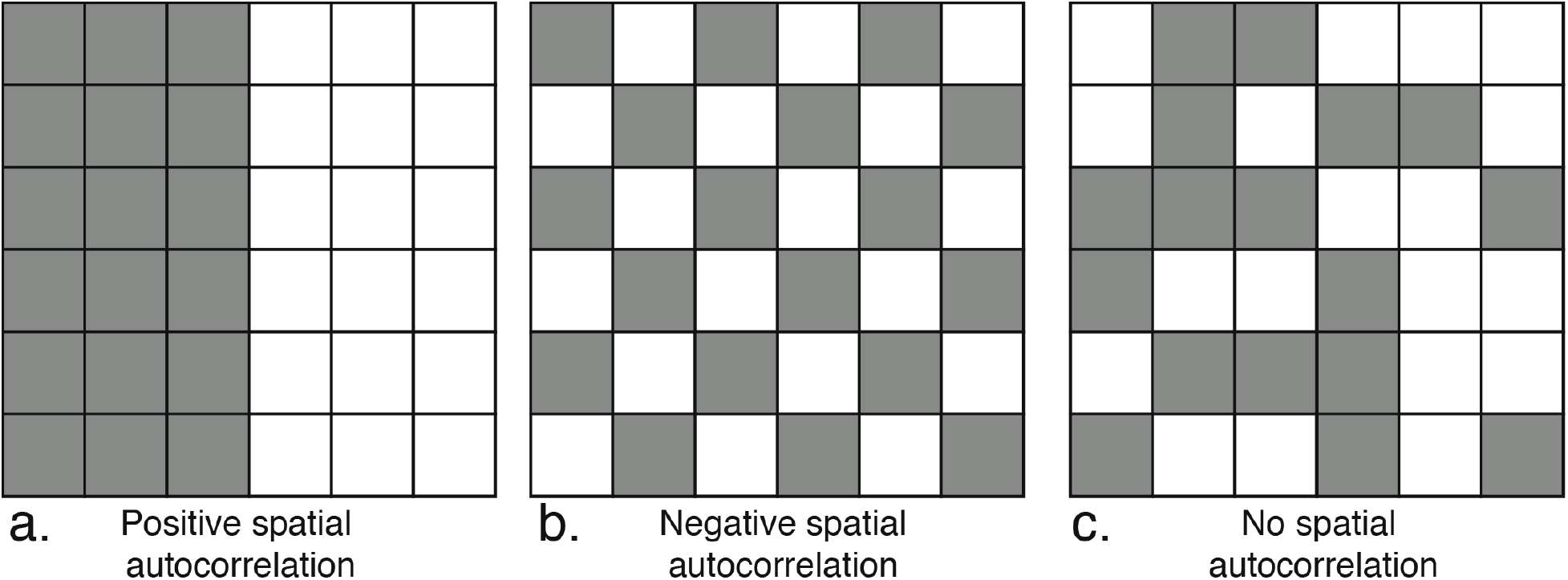
Conceptual models of spatial autocorrelation. Positive spatial autocorrelation (a) is associated with a clustering, or aggregating, phenomenon. Positive spatial autocorrelation was observed for all fungal and tree communities at small scales, and also across the extent of the study (Figure 2). Negative spatial autocorrelation (b) was observed for all fungal communities at intermediate scales (3.7–26 km). No spatial autocorrelation (c) is associated with randomly spaced distribution patterns for fungal communities and is also the null hypothesis used in the Mantel tests.

The sharp inversion of distance-decay patterns at intermediate distances (3.7–26 km) was unexpected given our hypotheses. Mantel tests revealed a pattern of negative spatial autocorrelation at the island scale (3.7–26 km; Table 1), indicating that, at this scale, fungal communities exhibit some form of spatial patterning. The classic example of negative spatial autocorrelation is often presented as a single species occurring in a checkerboard or interdigitating distribution pattern (Figure 4b). While it is difficult to interpret a multivariate dataset, like fungal community composition, on a checkerboard distribution, the negative spatial autocorrelation observed in this study does indicate a pattern of recurrent fungal community composition at intermediate scales. This anomalous inversion of the DDS trend is an example of Simpson’s paradox, where observations at the island scale do not adhere to trends seen in the whole dataset. This same paradox was observed in a different study of continental tropical plants where, at the smallest distance interval (0–600 km) and across the entire study (0–1600 km), plants exhibited positive spatial autocorrelation but at the two intermediate geographic distances (800–1100 and 1200–1600 km) the trend inverted and plant similarity increased with geographic distance [44]. For the larger distance classes of that plant study, environmental factors were more important for explaining plant community structure than was geographic distance. We observed a similar pattern in our study for belowground fungi. At smaller scales, geographic distance was the more explanatory factor for fungal community dissimilarity, whereas at larger scales (~ 100 km), environmental factors (i.e., tree community) were more important for explaining soil fungal community structure (Figure 3; Soil).

Spatial autocorrelation of fungal communities at a regional scale (between islands 80– 110 km) differed depending on the habitat from which fungi were sampled (Figure 2). Aboveground, *Selaginella* fungi demonstrated a distance decay slope consistent with trends across the entire study; *Selaginella* fungal community composition became more dissimilar with geographic distance. By contrast, soil fungal communities showed an inverted trend. Soil communities became increasingly similar with growing distance, a trend that was unexpected and implies factors other than geographic distance are structuring soil communities. Indeed, at large geographic scales (between islands 80–110 km), tree community composition explained more of the variability in soil fungal community composition than geographic distance (Figure 3; Soil), a hypothesis that had been previously demonstrated in ectomycorrhizal fungi of continental Europe [45].

The implication of this result is that exogenous factors, such as vegetation composition on separate islands (in this case, Tanna and Aneityum) play a larger role in structuring soil fungal communities than do endogenous factors, such as dispersal limitation. At the largest scale (80–110 km), fungal communities isolated from Soil and *Selaginella* showed opposite patters of spatial autocorrelation. In previous research of culture-based fungal endophytes, aboveground endophytes exhibited a distance-decay relationship at the regional scale but belowground endophytes did not [46]. It is unclear why fungal communities from different habitats would present such variable results at a large scale. Martiny et al. [24] found that for bacteria, the relative importance of different environmental parameters varied by scale, such that moisture was important at the local scale but temperature and nitrate concentrations were important at regional and continental scales. Since variation in environmental parameters is also scale-dependent we might expect habitat specific parameters, like edaphic characteristics, to exert differential influence on below- and aboveground fungal communities. A second possible explanation for the observed above/belowground spatial-autocorrelation paradigm may be related to the physical spacing of habitats. Soil, as a habitat, forms a nearly continuous habitat coextensive with the forest. Whereas *Selaginella,* as a habitat, is composed of individual plants dispersed in various patterns across the same landscape. Perhaps *Selaginella* acts more like islands compared to soil, resulting in a patchy habitat across the landscape.

### The relationship between plant and fungal communities

The effects of distance decay have long been recognized in plants [17] and animals [49], and efforts have been made to relate biogeographic patterns of plants and animals with those of microbes [12, 13]. In this study, we show that tree communities and fungal communities share a distance-decay relationship, indicating a strong relationship between fungal and tree communities. Furthermore, a direct comparison of plant and fungal DDS demonstrates that the spatial turnover of trees is greater than for above- and belowground fungi (Figure 2), a result previously reported for global soil fungi [10].

Past studies have indicated that host-plant species identity drives an association with specific foliar fungal communities [50, 51], suggesting that beta diversity of plant and endophyte fungal communities is linked. Our results indicate similar patterns (Table 2), but in this case, geographic distance is consistently a stronger factor in structuring aboveground fungal communities (Figure 3; Understory, *Selaginella).* By contrast, at large geographic scales, tree community composition was better at explaining soil fungi (Figure 3; Soil). This leads us towards two non-mutually exclusive hypotheses. First, at regional scales, correlations between plant and soil fungal communities are best explained by their similar responses to climatic and edaphic variables [10], whereas dispersal limitation is more important in structuring aboveground fungi.

Secondly, belowground plant-fungal symbioses are more important at structuring forest communities than are aboveground plant-fungal symbioses.

## Conclusion

Understanding how environmental factors and those associated with fungal biology contribute to the spatial dynamics of fungal communities is challenging. Here, we attempted to understand the spatial dynamics of above- and belowground fungi simultaneously, while also quantifying relationships with vegetation. Our results show that fungal communities of tropical-island forests exhibit strong spatial autocorrelation, and that the strength and type of autocorrelation is scale dependent. Fungal communities at local scales (< 50 m) are aggregated and show positive spatial autocorrelation, a trend that is inverted at the island scale (3.7–26 km). The spatial dynamics of fungal communities at a regional scale (~ 80–100 km) are more varied and are dependent on the habitat (belowground or aboveground) from which the fungal community is sampled. Across the extent of our study, geographic space is the dominating factor structuring aboveground fungal communities, whereas belowground, soil fungal community composition is better explained by forest community beta diversity. These results emphasize the importance of considering scale and spatial autocorrelation in analyses of fungal communities. Furthermore, our findings support a growing body of evidence supporting the idea that fungi and plants exhibit similar biogeographic patterns.

## Supporting information

Sampling methods

Fungal Diversity

Community x Community Mantel test

## Declarations

### *Availability of data and material* (data transparency)

The sequencing dataset analyzed during the current study is available in the NCBI Sequence Read Archive. (BioProject ID PRJNA634909)

Tree community, and transect data are available on Figshare (doi 10.6084/m9.figshare.12367475)

## Acknowledgments

The authors would like to thank the reviewers and Drs. Nicole Hynson, Tom Ranker, Nhu Nguyen, and Michael Kantar for improving this manuscript. We are also very appreciative of Presley Dovo and the Vanuatu Department of Forestry for logistical support. This project would not be possible without field support from the ever-growing network of people associated with *Plants mo Pipol blong Vanuatu.* We would also like to recognize the contributions of the late Philemon Ala who had been helping with *Plants mo Pipol* since its inception. Finally, we are grateful to the many communities of Aneityum and Tanna for their kindness, hospitality and for sharing so much invaluable knowledge, *tankyu tumas*.

---

**ESM_1.**
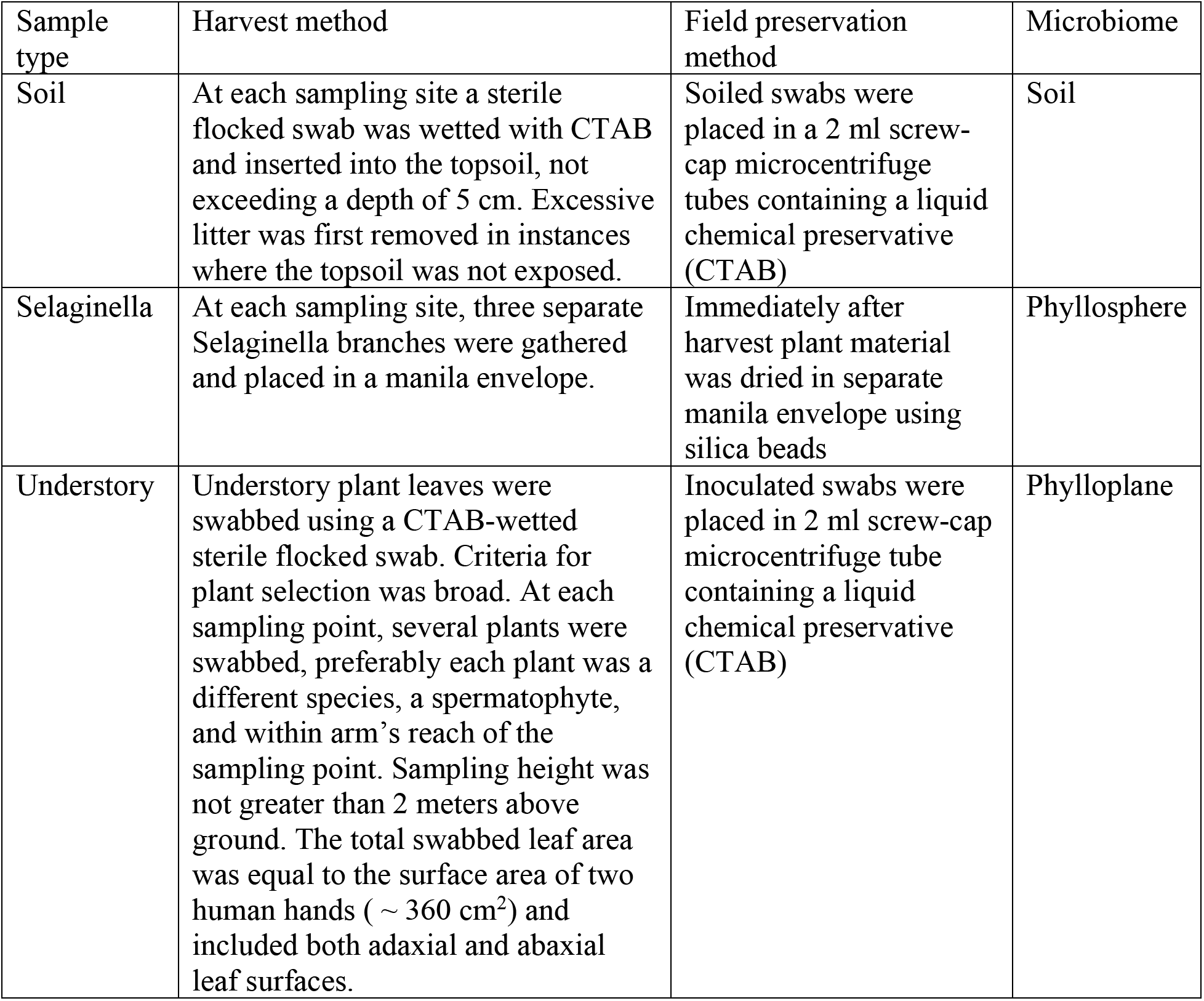
Fungal communities were sampled from three habitats Soil, Selaginella, and Understory (foliar epiphytes). Alongside are details of the harvesting and preservation method for each fungal community sample type. The chemical preservative was a CTAB solution (1M Tris-HCl pH 8, 5 M NaCl, 0.5 M EDTA and 20 g cetyltrimethylammonium bromide).

**ESM_2.**
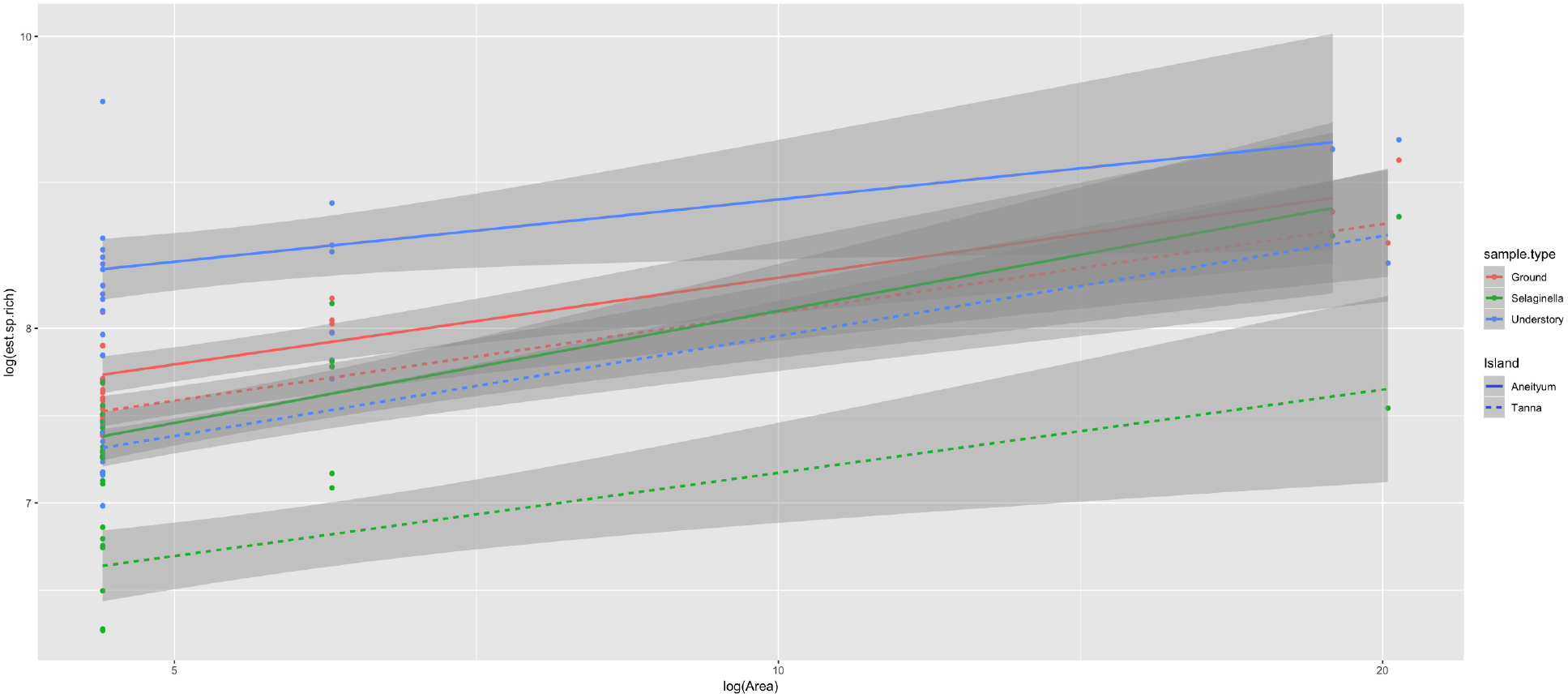
Species area curves (log-log transformed) for observed fungal OTUs parsed by island (Tanna (dotted) or Aneityum(solid)) and sample habitat (Soil (red), Selaginella (green), Understory Epiphytes (blue)). Important variability in observed species richness exists between islands, fungal habitat, and small areas. The shaded areas show the 95% confidence interval of the fit.

**ESM_3.**
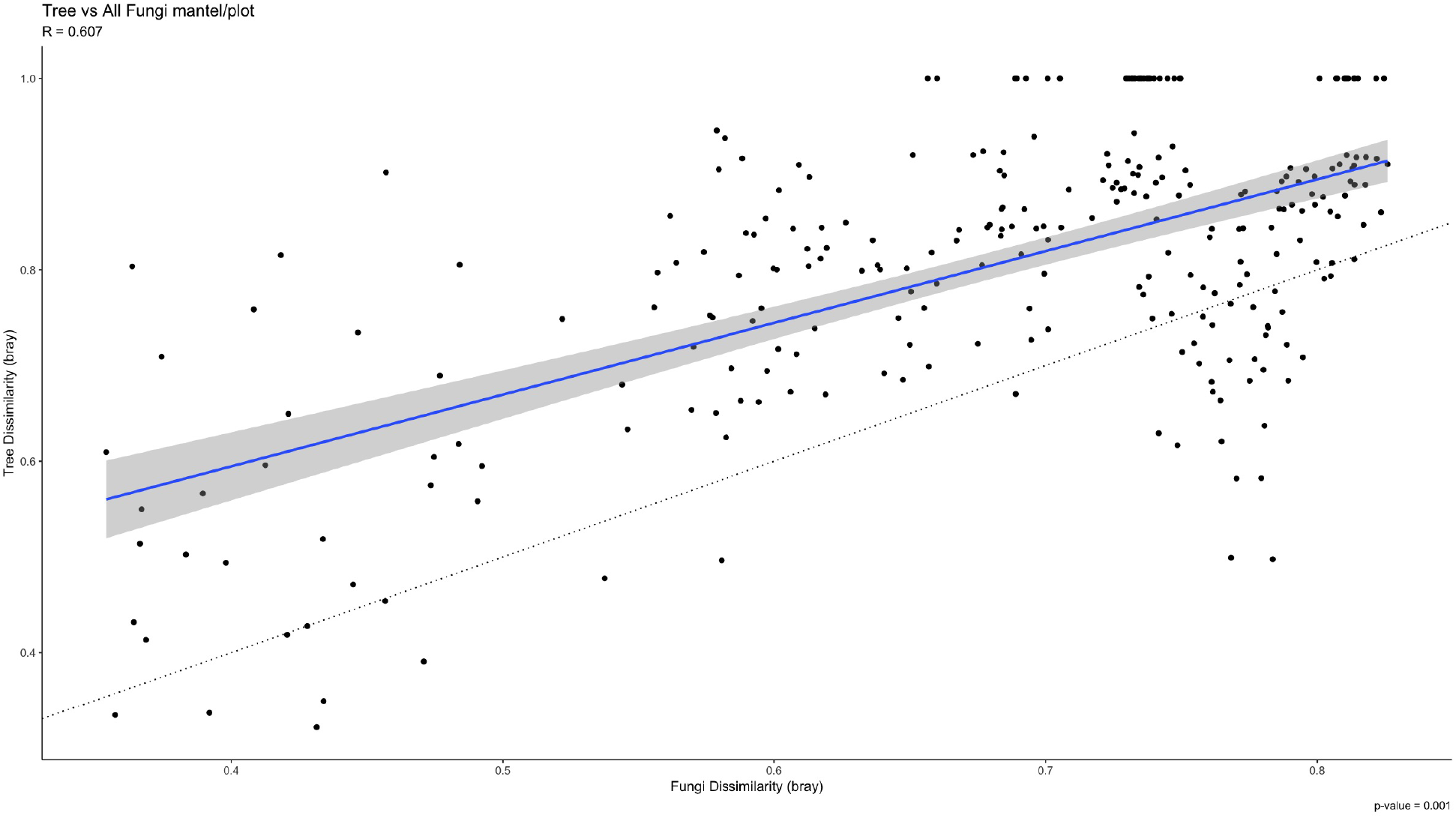
A partial Mantel test coupling fungal and tree Bray-Curtis community dissimilarities across the Vanuatu (n = 24 sites). This partial Mantel regression is accounting for collinearity of geographic distance using a third matrix of geographic distance. The linear regression of the pairwise Bray–Curtis distances for fungal and plant communities shows a significant positive correlation where increasingly similar plant communities are coupled with increasingly similar fungal communities (Mantel r = 0.607; p = 0.001). The shaded areas show the 95% confidence interval of the fit. Dotted line indicates a 1:1 slope for reference. Divergence from a slope of 1 indicates asynchronous betadiversity turnover.

