## Supplementary material for "Scale-dependent influences of distance and vegetation on the composition of aboveground and belowground tropical fungal communities": Sampling methods

ESM_1

*Fungal communities were sampled from three habitats: soil, phyllosphere (Selaginella), and phylloplane (understory epiphytes). Alongside are details of the harvesting and preservation method for each fungal community sample type. The chemical preservative was a CTAB solution (1 M Tris-HCl pH 8, 5 M NaCl, 0.5 M EDTA and 20 g cetyltrimethylammonium bromide).*

| Sample type | Harvest method | Field preservation method | Microbiome |
|---|---|---|---|
| Soil | At each sampling site a sterile flocked swab was wetted with CTAB and inserted into the topsoil, not exceeding a depth of 5 cm. Excessive litter was first removed in instances where the topsoil was not exposed. | Soiled swabs were placed in a 2 ml screw-cap microcentrifuge tubes containing a liquid chemical preservative (CTAB) | Soil |
| Selaginella | At each sampling site, three separate Selaginella branches were gathered and placed in a manila envelope. | Immediately after harvest plant material was dried in separate manila envelope using silica beads | Phyllosphere |
| Understory | Understory plant leaves were swabbed using a CTAB-wetted sterile flocked swab. Criteria for plant selection was broad. At each sampling point, several plants were swabbed, preferably each plant was a different species, a spermatophyte, and within arm's reach of the sampling point. Sampling height was not greater than 2 meters above ground. The total swabbed leaf area was equal to the surface area of two human hands ( ~ 360 $cm^2$) and included both adaxial and abaxial leaf surfaces. | Inoculated swabs were placed in 2 ml screw-cap microcentrifuge tube containing a liquid chemical preservative (CTAB) | Phylloplane |
