## Supplementary material for "Scale-dependent influences of distance and vegetation on the composition of aboveground and belowground tropical fungal communities": Fungal Diversity

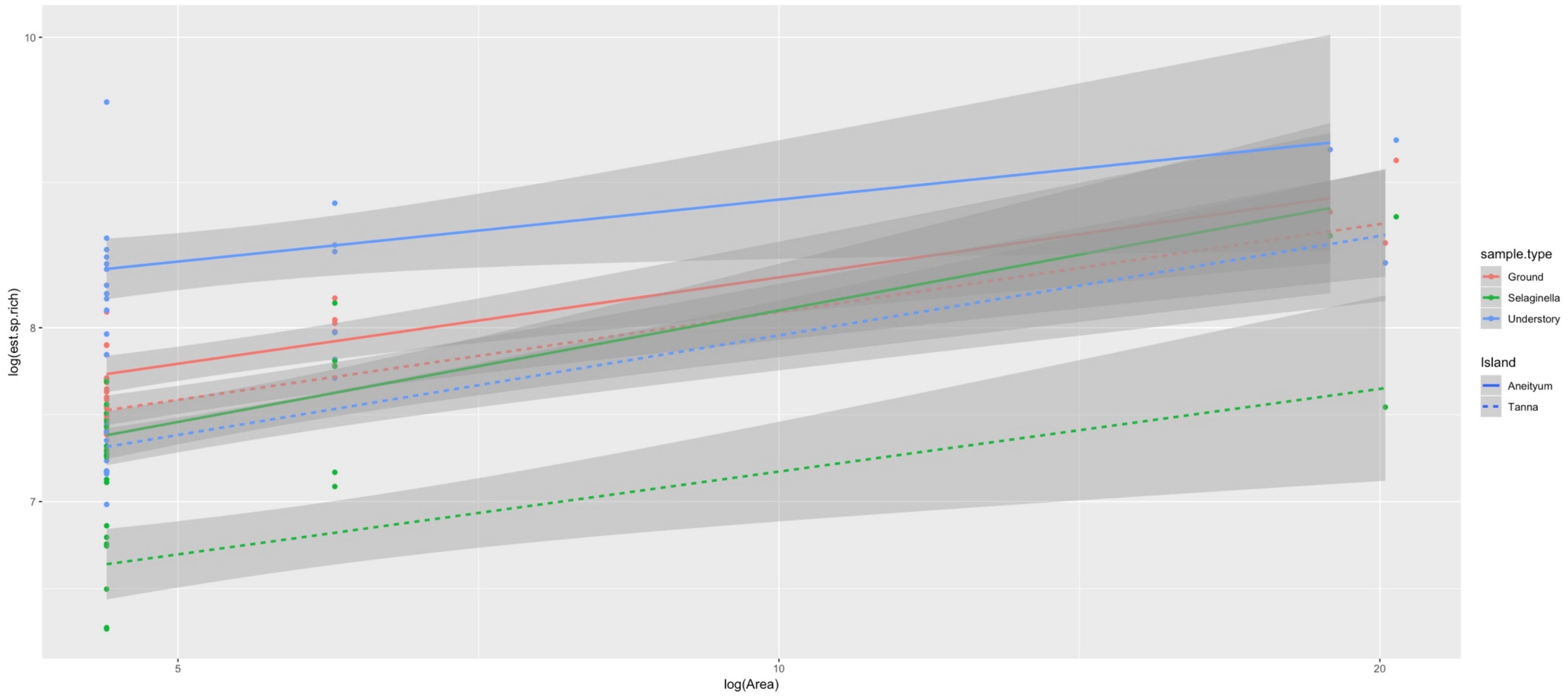
